# Step-wise mechanical unfolding and dissociation of the GAIN domains of ADGRG1/GPR56, ADGRL1/Latrophilin-1 and ADGRB3/BAI3: insights into the mechanical activation hypothesis of adhesion G protein-coupled receptors

**DOI:** 10.1101/2023.03.14.532526

**Authors:** Chaoyu Fu, Wenmao Huang, Qingnan Tang, Minghui Niu, Shiwen Guo, Tobias Langenhan, Gaojie Song, Jie Yan

## Abstract

Adhesion G protein-coupled receptors (aGPCRs) are a large family within the superfamily of G protein-coupled receptors involved in various physiological processes. One unique feature of aGPCRs is their long N-terminal extracellular regions (ECRs), which contain adhesive domains and a GPCR autoproteolysis-inducing (GAIN) domain. This GAIN domain promotes autoproteolytic cleavage of aGPCRs into N- and C-terminal fragments (NTF, CTF, respectively) after receptor biosynthesis. aGPCR signaling involves an interplay between the NTF and CTF that can be mechanically activated or modulated. However, how force affects the conformation/structure of the GAIN domain as a central structural element in aGPCR activation remains largely unknown. In this study, we investigated the mechanical stability of the GAIN domains of three aGPCRs from subfamilies B, G and L at a loading rate of 1 pN/s. Our findings demonstrate that the GAIN domains can be destabilized by forces from a few to 20 piconewtons (pN). Specifically, for the autocleaved aGPCRs, ADGRG1/GPR56 and ADGRL1/Latrophilin-1, forces over this range can cause detachment of the GAIN domain from the membrane-proximal Stachel element, which serves as an endogenous tethered agonist to aGPCRs, typically preceded with GAIN domain unfolding. For the non-cleavable aGPCR ADGRB3/BAI3, the GAIN domain undergoes complex mechanical unfolding over a similar force range. We also demonstrate that detachment of the GAIN domain can take place during cell migration, provided that the linkage between aGPCR and extracellular matrix is sufficiently stable. These results suggest that both structural stability of the GAIN domain and NTF/CTF dissociation are sensitive to physiological ranges of tensile forces, providing insights into the mechanical activation hypothesis of aGPCRs.

## I. INTRODUCTION

G protein-coupled receptors (GPCRs) are a superfamily of transmembrane receptors characterized by seven transmembrane (7TM) *α*-helices. The majority of GPCRs are activated by ligand binding, which triggers signal transduction resulting in cellular responses. Extracellular ligand molecules binding to GPCRs causes a conformational change in the receptor’s extracellular region or 7TM domain, which is then allosterically propagated across the plasma membrane. Ultimately, this is brought about by a conformational change in the receptor’s transmembrane region. This process is critical for cellular signaling and has been extensively studied [1].

GPCRs can be classified into five major families based on the phylogenetic grouping of their 7TM helices: glutamate, rhodopsin, adhesion, frizzled, and secretin [2–5]. Among these families, the adhesion GPCRs (aGPCRs) constitute the second largest family with 33 mammalian homologs [2]. Most aGPCRs possess a large multidomain extracellular N-terminal region (ECR) that facilitates adhesive interactions with the extracellular matrix (ECM) or membrane proteins on neighboring cells [6–9]. Thirty-two out of the 33 mammalian aGPCRs contain a conserved GPCR autoproteolysis-inducing (GAIN) domain that harbors a GPCR proteolysis site (GPS), which is embedded within the GAIN domain [8, 10, 11]. The GAIN domain contains a tethered agonist element (TA, Stachel) as part of the GAIN fold, which is necessary and sufficient for the activation of aGPCRs [12, 13].

In some aGPCRs, the GAIN domain is capable of undergoing autoproteolysis, resulting in the formation of two non-covalently associated fragments: the N-terminal fragment (NTF), and the rest of the receptor, which is referred to as the C-terminal fragment (CTF) [11, 14–17]. The CTF consists of the TA that is connected to the first transmembrane helix of the 7TM domain [8, 12, 13]. However, self-cleavage is not a general feature of aGPCRs as the GAIN domains of a few aGPCRs are incapable of self-proteolysis (see review in [9, 18]).

aGPCRs play critical roles in a wide variety of biological processes, including neuronal development, immune cell function, and cancer progression [19–23]. Despite the important functions of aGPCRs, the activation mechanisms is still incompletely understood. aGPCR activation requires an endogenous tethered agonist (Stachel), which is part of the GAIN domain [12, 13]. Stachel exposure from its encasement within the GAIN domain is currently supported by two models. The dissociation (one-and-done) model requires the physical disruption of the NTF/CTF heterodimer [12, 24–26], while the non-dissociation (tunable) model posits that Stachel-7TM domain engagement occurs through partial allosteric movements of the GAIN domain in intact NTF/CTF aGPCR heterodimers [10, 13, 17, 27, 28]. Both scenarios have received support from recent structural studies of active aGPCR 7TM domains [29–35]. The existence of self-cleavable and non-cleavable aGPCRs underline the feasibility of both models, complicating pharmacological advances.

Several self-cleavable [25, 36–40] and non-cleavable [41] aGPCRs show sensitivity to mechanical stimuli applied to their NTFs, indicating mechanical activation as an adequate mode of activation. However, the exact mechanism of this mechanical activation process remains unknown. A mechanical activation model has been proposed, where force-induced NTF/CTF dissociation exposes the Stachel to activate aGPCRs [42–45]. Similarly, mechanical activation can also explain non-dissociative aGPCR activation, where forces applied to NTFs may allosterically change the GAIN domains conformation and transiently expose the Stachel [17, 41].

While the mechanisms of mechanical activation are both interesting and important, it is still unclear whether the typical physiological range of forces, which vary from a few pN to tens of pN [46, 47], can induce NTF/CTF dissociation or significant conformational changes of the GAIN domains. To bridge this knowledge gap between mechanical activation models of aGPCRs and their physiological functions, we investigated the force-response of the GAIN domains of three representative aGPCRs, including ADGRG1/GPR56 and ADGRL1/LPHN1, which both possess cleavability but different GAIN domain organizations, as well as ADGRB3/BAI3, which has a non-cleavable GAIN domain. We applied forces to individual GAIN domains of these aGPCRs and recorded their force-dependent dissociation/conformational changes in real-time at a nanometer (nm) resolution using an inhouse constructed magnetic-tweezers setup [48, 49]. We also investigated the effect of cell migration on the dissociation of the NTF/CTF of GPR56 using a surface-bound artificial ligand that forms a covalent bond with the N-terminus of GPR56.

Our results reveal that 1) the subdomains in the GAIN domain undergo structural unfolding within a few piconewtons (pN) of forces, 2) for aGPCRs with cleavable GAIN domain, the physiological range of forces can cause NTF dissociation in addition to domain unfolding, and 3) the NTF/CTF dissociation could occur during cell migration when the aGPCR-extracellular matrix (ECM) linkage is sufficiently stable. Our results provide a physical basis that sheds light on the two mechanical activation models, and how cell migration could play a role in the acivation of the receptors.

## II. RESULTS

### A. Dissociation of GPR56 and LPHN1 during force loading

Figure 1a shows schematic illustrations of the experimental design and the protein constructs for GPR56, LPHN1, and BAI3 used in this study. The cleavable GPR56 construct contains a cleavable GAIN domain, with two *α*-helices in its GAIN A sub-domain [50]. The cleavable LPHN1 construct contains a GAIN domain consisting of five-helix GAIN A subdomain and a GAIN B with similar corresponding structures to those in the non-cleavable BAI3 [11]. BAI3 construct includes an additional HormR domain at the N-terminus.

**Fig. 1.**
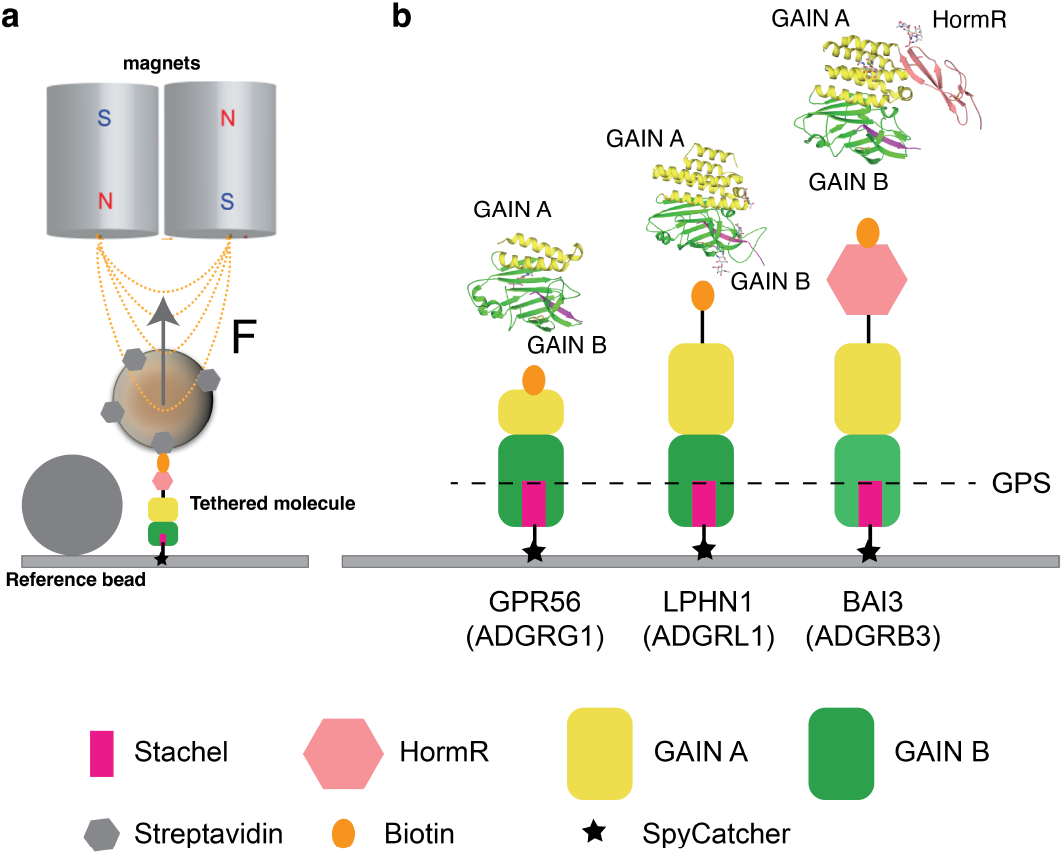
System Setup and Tethered Configuration of GPR56, LPHN1, and BAI3. (a) An aGPCR’s GAIN domain tethered between a 2.8-*μ*m-diameter superparamagenetic bead is subjected to an external force excerted by a pair of permanent magnets. (b) The GAIN domains of GPR56, LPHN1, and BAI3 are depicted in the illustration. The GPS are located within the GAIN B subdomains.

Each protein construct has a biotinylated AVI tag at the N-terminus and a SpyTag at the C-terminus. The C-terminus is immobilized to a SpyCatcher-coated coverslip surface [51], while the N-terminus is linked to a 2.8-*μ*m-diameter super-paramagnetic bead (Fig. 1). Tensile forces are applied to individual protein tethers using an in-house constructed magnetic tweezers setup [48, 49], by attaching one end of the superparamagnetic bead to a pair of permanent magnets placed above the chamber. The magnitude of the force is controlled by adjusting the magnet-bead distance, and the change in bead height from the coverslip surface is recorded at nanometer resolution with a sampling rate of 200 Hz. Details of the force calibration, bead height determination, and force level control are available in our previous publications [48, 49].

We began our investigation by examining the mechanical response of the GPR56’s GAIN domain to an increasing force at a loading rate of 1.0 ± 0.1 pN/s. Figure 2a displays the representative changes in the height of beads from four independent tethers. The curves are shifted along the *Y*-axis for better visualization. We note that while the change in bead height is a result of both bead rotation and extension change of the molecule during force change, the bead height change during a stepwise change is equivalent to the molecular extension change [49].

**Fig. 2.**
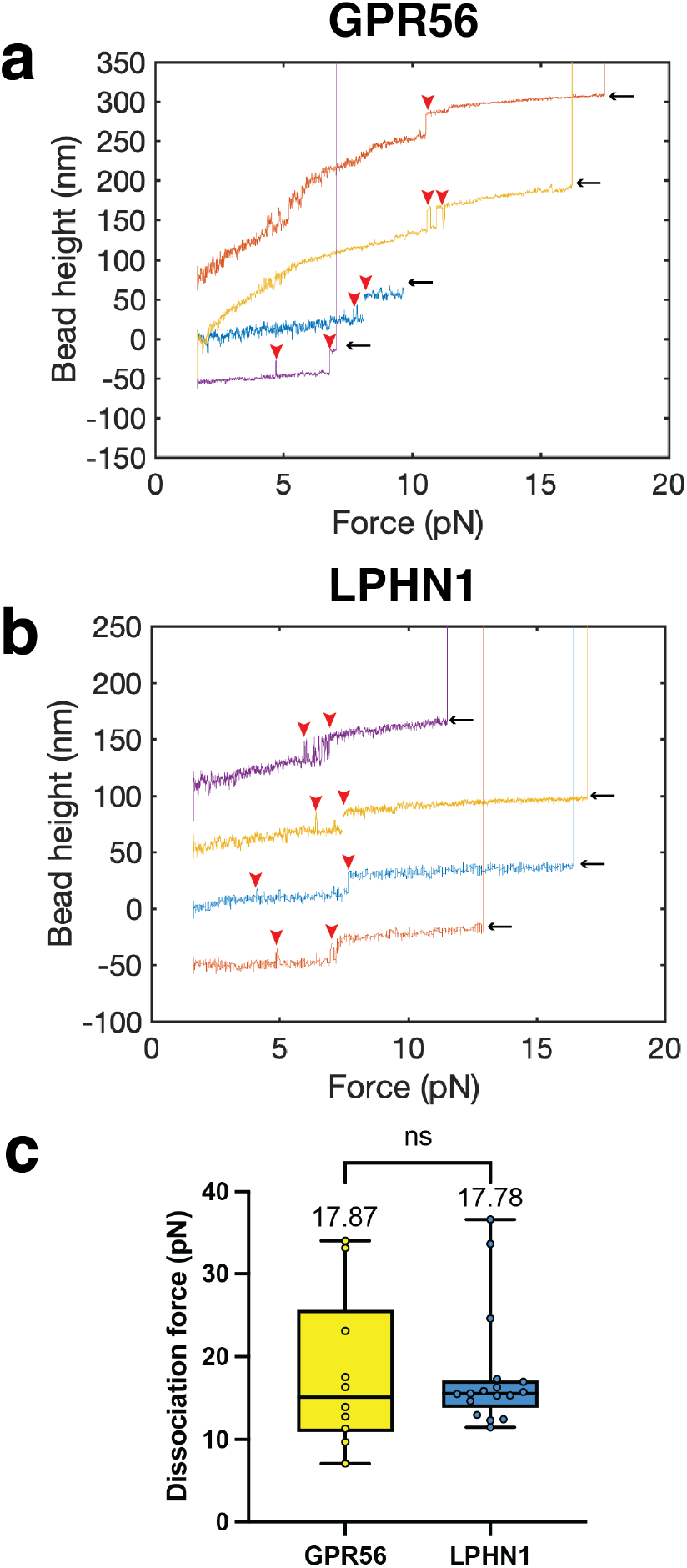
NTF/CTF dissociation of the GAIN domains of GPR56 and LPHN1. (a) Representative force-bead height curves of four tethers for GPR56’s GAIN domain during force loading until NTF/CTF dissociation. Black arrow: NTF-CTF dissociation event; Red arrowhead: GAIN domain unfolding event. (b) Representative force-bead height curves of four independent tethers for LPHN1’s GAIN domain during force loading until NTF/CTF dissociation. Black arrow: NTF-CTF dissociation event; Red arrowhead: GAIN domain unfolding event. (c) Box plots of forces where NTF/CTF dissociation events were observed, which provides information of the medians and IQRs. The means are indicated. A fixed loading rate of 1.0 ± 0.1 pN/s was applied.

At this loading rate, the time traces reveal structural transitions at forces between 5-10 pN, indicated by stepwise extension changes (Figure 2a). The tethers were broken when forces were further increased. The same type of experiments conducted for LPHN1 revealed similar mechanical responses over a similar force range (Figure 2b).

Figure 2c summarizes the NTF/CTF dissociation forces obtained for more than 10 tethers for each construct. The forces were found distributed with interquartile range (IQR) from 10-20 pN for the GPR56’s GAIN domain and 14-17 pN for LPHN1’s GAIN domain. These results suggest that the NTF/CTF dissociation for both GPR56 and LPHN1 occurs within a force range of 10-20 pN at our loading rate.

### B. Force-dependent GAIN domain structural changes of GPR56 and LPHN1

As shown in Figure 2, extensive structural changes of the GAIN of GPR56 and LPHN1 occurred at forces before NTF/CTF dissociation, indicated by stepwise extension increases (unfolding) during the force-loading process. These force-dependent unfolding and refolding events (more than 50) from more than 10 tethers are represented in force-stepsize graphs (Fig. 3a), the box plots of the unfolding forces (Fig. 3b) and the unfolding stepsizes (Fig. 3c).

**Fig. 3.**
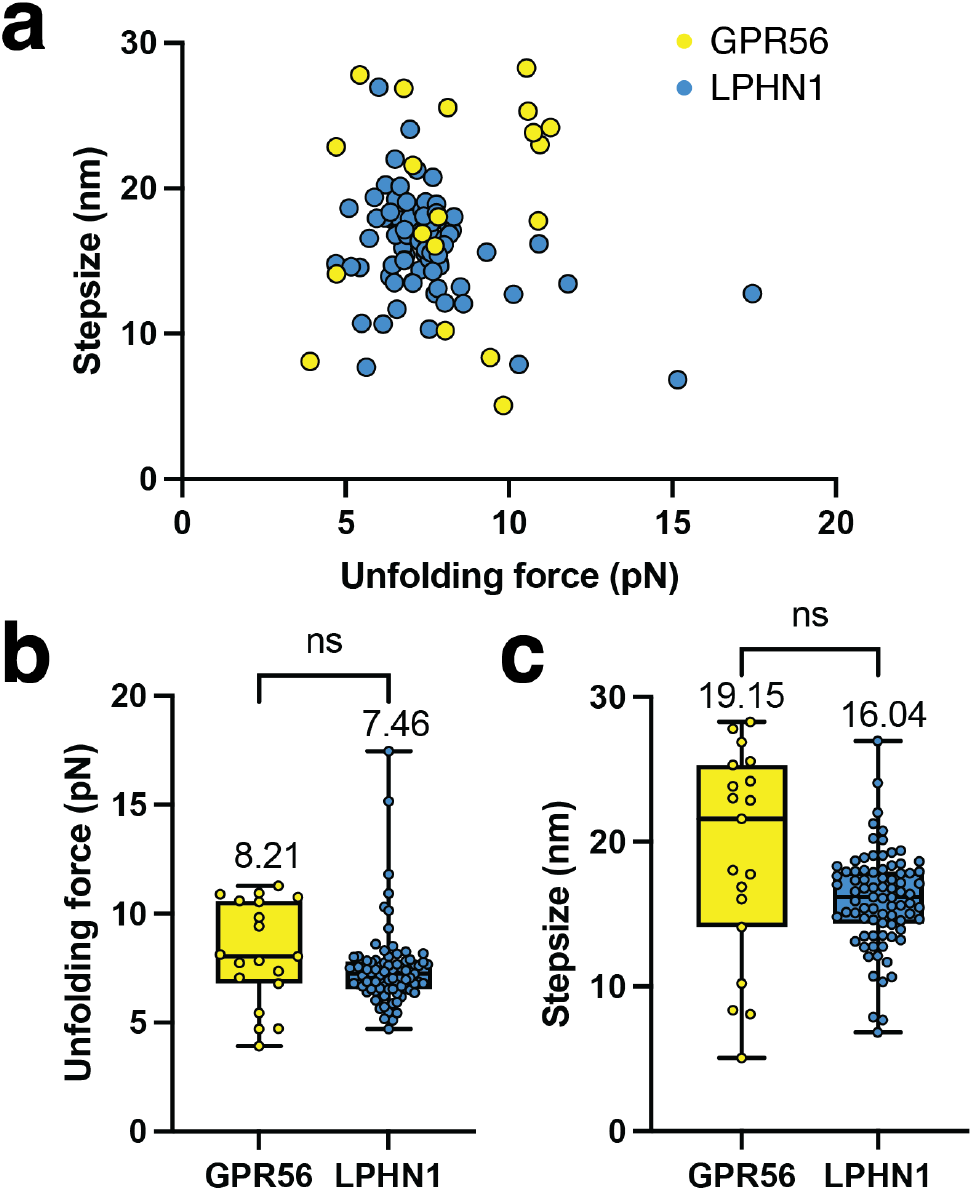
Force-induced unfolding of the GAIN domains of GRP56 and LPHN1. (a) Two-dimensional scatter plot depicting the force-step size data for the unfolding steps preceding the dissociation of NTF/CTF. (b-c) Box plots of forces (b) and stepsizes (c) where unfolding events were observed, which provides information ofthe medians and IQRs. The means are indicated. A fixed loading rate of 1.0 ± 0.1 pN/s was applied.

Based on the data, it appears that the majority of unfolding forces for both GAIN domains occur over a force range from 5 pN to 10 pN at a loading rate of 1 pN/s. The stepsizes mainly range from 10-30 nm. Notably, the domain unfolding forces are smaller than the NTF/CTF dissociation forces, as depicted in Figure 2c.

### C. Mechanical responses of BAI3 GAIN domain during force loading

In contrast to the GAIN domains of GPR56 and LPHN1, the GAIN domain of BAI3 is not cleavable. As a result, our examination of the BAI3 GAIN domain focused on force-dependent conformational changes. Figure 4a displays bead height - force curves obtained from a single representative tethered BAI3 construct subjected to multiple consecutive force-loading processes. Following each loading, the force was reduced to 1 pN for 1 minute to allow the domains to refold.

**Fig. 4.**
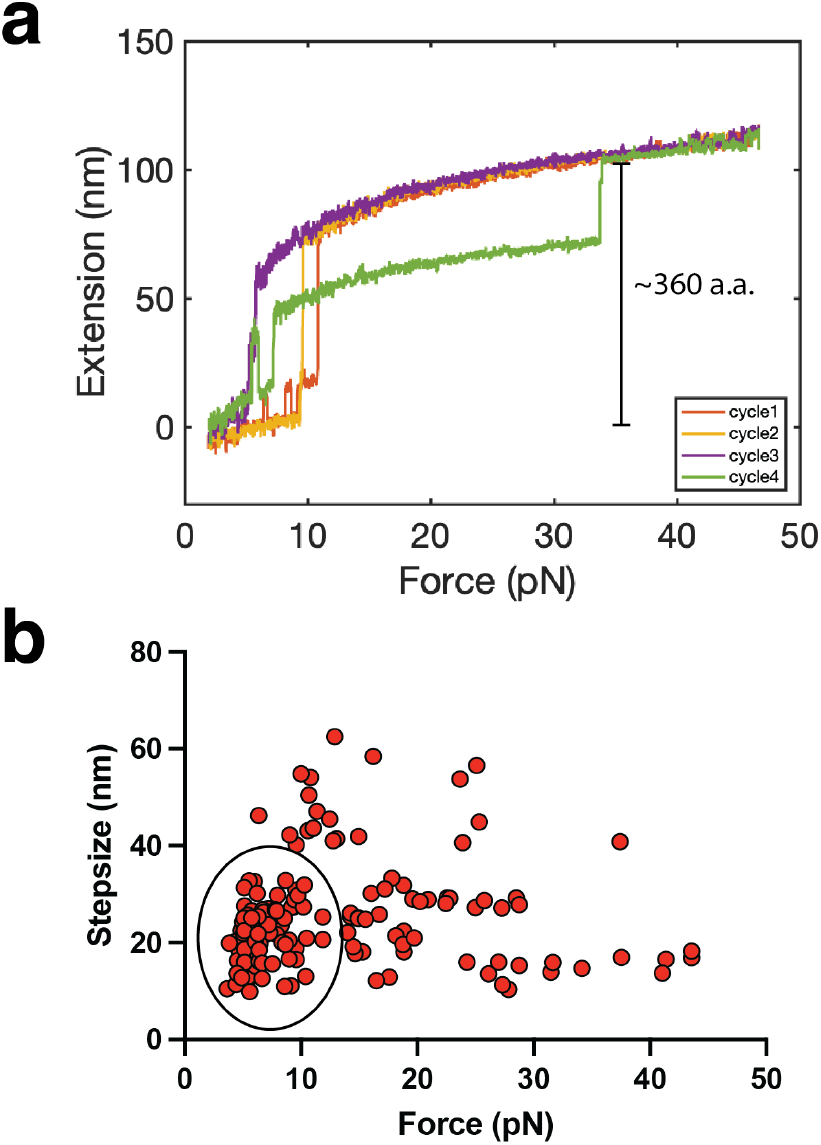
Force-induced unfolding of the GAIN domain of BAI3. a) The panel show representative bead height-force curves obtained from successive force loading processes at a loading rate of 1.0 ± 0.1 pN/s. Multiple steps of unfolding events were observed in each force loading process. (b) The panel shows a twodimensional scatter plot of the force-step size data for the unfolding events. Data in the circle represent unfolding events that are similar to those observed for the GAIN domains of GPR56 and LPHN1 preceding the NTF/CTF dissociation.

The curves show that there are multiple stages of unfolding signals during each force loading process. When the force exceeds 30 pN, the curves overlap, indicating complete unfolding of all the structural elements within the GAIN domains at that level of force. Based on the force-dependent step sizes and assumeing a typical bending persistence of 0.8 nm for low-force response of polypeptide polymers [48, 52], it is estimated that tension releases more than 350 amino acids of polypeptide after unfolding. This suggests that the GAIN and HormR domains of BAI3, which comprise around 360 amino acid residues, completely unfolds at forces within 30 pN.

The presence of multiple steps at widely spread forces in each force loading process suggests differential mechanical stabilities of the subdomains in the BAI3 GAIN domain. Additionally, the presence of unfolding steps in each force loading process indicates that the unfolded GAIN domain can quickly refold at lower forces.

Figure 4b shows the force-stepsize graph for the unfolding signals of the GAIN domain of BAI3 obtained from over 10 tethers. Interestingly, while the data is spread out in both unfolding forces and stepsizes, there exists a clear circled cluster of data within a force range of 5-10 pN and stepsizes ranging from 10 - 30 nm. These results are similar to those observed for the GAIN domains of GPR56 and LPHN1 before the NTF/CTF dissociation.

### D. Direct visualization of GAIN domain cleavage during cell migration

The NTFs of aGPCRs contain domains that are capable of associating with ECM or membrane receptors on other cells. This association is necessary to establish mechanical stretching of aGPCRs. If these NTF-mediated physical linkages are sufficiently strong under mechanical stretching, the aGPCRs adhered to the ECM may reach a tension level above the pN range, which could cause conformational changes in the GAIN domain and result in NTF/CTF dissociation during cell migration.

We tested this hypothesis by a cell-based assay that allows for direct visualization of the dissociation of NTF and CTF of GPR56 during cell migration. To achieve this, we used an artificial covalent ligand that forms a stable linkage between NTF and ECM.

To carry out the experiment, we created two GPR56 constructs: Spy-GPR56-WT-GFP and Spy-GPR56-T383A-GFP. We inserted a SpyTag between the signal peptide and the pentraxin and laminin/neurexin/sex hormone-binding globulin-like (PLL) domain and added a GFP tag after the 7TM domains in both constructs (Figures 5a). The Spy-GPR56-T383A-GFP was a non-cleavable mutant that lacked the ability to undergo GAIN domain cleavage [53]. On the other hand, Spy-GPR56-WT-GFP was cleavable. We confirmed the cleavage ability of Spy-GPR56-WT-GFP and the inability of Spy-GPR56-T383A-GFP to undergo GAIN domain cleavage through independent Western blot analysis (Figures 5b).

**Fig. 5.**
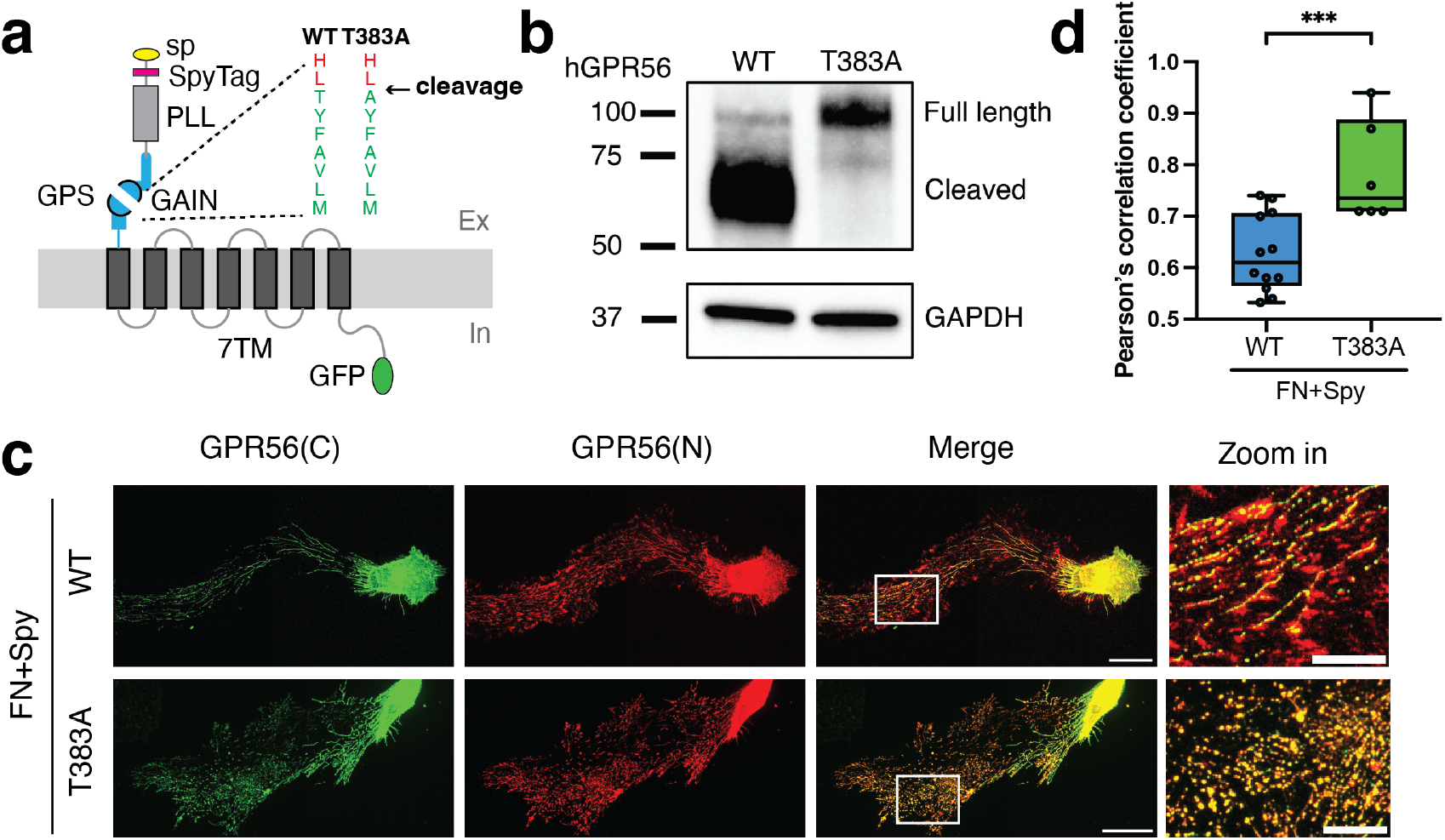
GPR56 NTF/CTF dissociation during cell migration. Direct observation of GAIN domain dissociation during cell migration. a. Schematic of GPR56 fusion protein used in the study. The extracellular domain of GPR56 consists of a signal peptide (sp), a Pentraxin/Laminin/neurexin/sex-hormone-binding-globulin-Like (PLL), and a GAIN domain. A SpyTag was introduced between the signal peptide and the PLL domain. The GAIN domain is cleaved between amino acids 382 and 383 at a conserved GPS. A non-cleavable mutant of GPR56 (Spy-GPR56-T383A-GFP) was created suppressing cleavage of the GAIN domain. b. Western blot of whole-cell lysates from HFF cells expressing either the wild-type GPR56 construct (Spy-GPR56-WT-GFP) or the non-cleavable construct (Spy-GPR56-T383A-GFP). c. HFF cells were transfected with either Spy-GPR56-WT-GFP or Spy-GPR56-T383A-GFP constructs, and then spread and migrated on surfaces coated with FN+Spy. NTF-CTF dissociation was measured by loss of colocalization of the C-terminal fragment, which is visualized by GFP (green), and N-terminal fragment, which is visualized by the staining of antibody targeting NTF (red). The loss of colocalization was observed in the cells expressing Spy-GPR56-WT-GFP but to a significantly lesser degree when GAIN domain self-cleavage was suppressed. Scale bar, 30*μ*m for merge, 10*μ*m for zoom-in. d. Pearson’s correlation coefficient between the N-terminal fragment (GPR56-N) and the C-terminal fragment (GPR56-C) of GPR56 in the conditions of c (N>5).

We transfected human foreskin fibroblast (HFF) cells, which do not express endogenous GPR56, with either Spy-GPR56-WT-GFP or Spy-GPR56-T383A-GFP. The cells were then seeded onto fibronectin mixed with Spy-Catcher, referred to as FN+Spy surface, to provide an integrin-based substrate for cell migration. SpyCatcher acted as an artificial ligand that could covalently associate with the SpyTag on the NTFs of Spy-GPR56-WT-GFP and Spy-GPR56-T383A-GFP. The resulting covalent bond between SpyCatcher and SpyTag allowed the cleaved NTFs to be retained on the surface for later immunostaining analysis.

Our observations indicate that Spy-GPR56-WT-GFP was clearly cleaved on the FN+Spy surface, as shown by the mis-colocalization of the receptor’s N-terminal and C-terminal regions. In contrast, cells expressing the non-cleavable Spy-GPR56-T383A-GFP exhibited well-co-localized N-terminal and C-terminal regions on the FN+Spy surfaces (Figures 5c). Pearson’s correlation coefficients supported these findings (Figures 5d). These results suggest that NTF/CTF dissociation may occur during cell migration if the NTF/CTF linkage is sufficiently stable. Our in vitro single-molecule studies also demonstrated that GAIN domain conformational changes typically precede NTF/CTF dissociation. Therefore, it is reasonable to suggest that cell migration may also lead to conformational changes in GAIN domains.

## III. MATERIALS AND METHOD

### Protein expression and purification

Codon-optimized cDNA sequence of human BAI3 fragment (508-873), LPHN1 fragment (533-851), and GPR56 fragment (173-394) were synthetized and inserted into a customized pLEXm vector with N-terminal signal peptide and C-terminal His6 tags, respectively. For each fragment, an avi-tag and a SPY tag were further added immediately to the N-terminus and C-terminus, respectively. HEK293F GnTI-cells was cultured with FreeStyle 293 Expression minute rotating (50mm amplitude shaker), 37 ^ř^C, 5% CO2, 95% humidity environment. When cultured to 1-2 million/mL, the cells were transiently transfected with DNA: polyethylenimine at 1:3 wt/wt. After 3 to 5 days, cell supernatants were harvested and incubated with Ni-NTA beads (Smart-Lifesciences, China) in 4 ^ř^C for about 2h. Beads were washed with 20 mM HEPES, pH 7.5, 500 mM NaCl, 30 mM imidazole and eluted with the similar buffer except imidazole reached to 300 mM. The proteins were further purified using size exclusion chromatography (SEC) with a Superdex S200 increase gel filtration column (cytiva) equilibrated with 20 mM HEPES, pH 7.5, 150 mM NaCl. After concentration, the proteins were further subjected to biotinylation using a bioA kit (AVIDITY) and checked with western blot. Mutants were acquired with quickchange site-directed mutagenesis kit and purified similarly. The optimal biotinylated samples were subjected to magnetic tweezer study.

### Single-molecule manipulation

The single-molecule manipulation experiments were carried out on a custom-built magnetic-tweezers setup [54] that record bead images at a 200 Hz sampling rate. For the given magnets and bead, the force is solely dependent on the distance between the magnet and the bead, *F*(*d*), which can be calibrated based on its fluctuation at low force, as described in previous publication, with 10% uncertainty due to the heterogeneous manufactured bead sizes [54]. The details of the chamber and microbeads preparation, sample preparation, force calibration and the loading rate control are described in our recent review paper [49].

### Cell culture and reagents

Human Foreskin Fibroblasts (HFF) cells were cultured at 37 ^ř^C in a 5% CO2 incubator in high-glucose Dulbecco’s Modified Eagle Medium (DMEM) supplemented with 10% Fetal Bovine Serum (FBS), 100IU/ml Penicillin/Streptomycin and 1 mM sodium pyruvate. The cells were tested for mycoplasma contamination and found negative. Antibodies: Anti-GPR56 Antibody (G-6) N-terminal (santa crutz, sc-390192).

Human GPR56 cDNA was cloned into a pCDNA3-EGFP vector. The forward primer contained a SpyTag introduced downstream of the signal peptide sequence in the cDNA. Mutant GPR56-T383A construct was created using primers 5-CAACCACTTGgCCTACTTTGC-3 and 5-CAGAAGCAGGATGTTTGG-3 by site-directed mutagenesis using Q5 ő Site-Directed Mutagenesis Kit (NEB). These mutant constructs were confirmed by Sanger sequencing. HFF cells were seeded into a 6-well plate with 60-70% confluence at day 0 and transfected with either Spy-GPR56-WT-GFP or Spy-GPR56-T383A-GFP using a Neon electroporation system (Invitrogen) following the manufacturer’s instructions on day 1. The transfected single HFF cells were seeded on the glass bottom dishes coated with Fibronectin mixed with SpyCatcher. After 12 h cell spreading, cells were fixed for immunofluorescence.

### Immunofluorescence

After undergoing cell spreading, HFF cells were fixed with a pre-warmed solution of 4% paraformaldehyde in PBS at 37 ^ř^C for 15 minutes. This was followed by permeabilization with 0.2% Triton X-100 in TBS for 30 minutes at room temperature to allow for antibody penetration. To minimize non-specific binding, the samples were then treated with a blocking solution of 1% Bovine Serum Albumin (BSA) in TBS for 1 hour. The samples were then incubated with primary antibodies overnight at 4 ^ř^C, followed by an incubation with secondary antibodies for 1 hour at room temperature.

### Western blot

Confluent cells were lysed in a RIPA buffer (Sigma) and a protease inhibitor (Roche) for 20 minutes at 4 ^ř^C. The amount of proteins extracted was determined using the BCA assay (Bio-Rad). The extracted proteins were separated by 4-20% SDS polyacrylamide gel electrophoresis (PAGE) and transferred to nitrocellulose membranes (Bio-Rad) using an electrotransfer process for 7 minutes (Trans-Blot Turbo system). The membranes were blocked with a solution of 5% Bovine Serum Albumin (BSA) in TBS 0.05% Tween 20 to prevent non-specific binding. The membranes were then incubated with primary antibodies (diluted 1:1000) overnight at 4 ^ř^C, followed by an incubation with secondary HRP antibodies (diluted 1:2000) for 1 hour at room temperature. The results were visualized using the ChemiDoc chemoluminescence detection system (Bio-Rad) and quantified using Image Lab. GAPDH was used as a reference protein to control for variations in protein loading.

### Statistical analyses

The cell migration measurements were performed at least three times in triplicate. Origin (OriginLab), MATLAB (The MathWorks), and Prism (GraphPad Software) were utilized for data analysis and graph plotting. Statistical significance was determined using Welch’s t-test. Additionally, the co-localization of GPR56-GFP with the NTF of GPR56 was analyzed by calculating the Pearson’s correlation coefficient between the two channels.

## IV. DISCUSSION

To summarize, we have investigated the mechanical responses of two autocleavable aGPCRs, GPR56 and LPHN1, and a non-cleavable aGPCR, BAI3, at a singlemolecule level using a magnetic-tweezers setup. We found that the GAIN domains of all three aGPCRs are sensitive to tensile forces, ranging from a few pN to about 20 pNs, at a physiologically relevant force-loading rate of 1 pN/s (see next paragraph). For GPR56 and LPHN1, NTF/CTF dissociation occurs at forces between 10-20 pN. Prior to the dissociation, we observed dynamic unfolding and refolding associated with large stepsizes of tens of nanometers, suggesting extensive unfolding of the GAIN domains. By using SpyCatcher as an artificial ligand, we clearly demonstrated the dissociation of SpyTag fused NTF from CTF in a cell migration assay. Taken together, these results demonstrate that the structural integrity of the GAIN domains of these aGPCRs is precisely regulated by pN forces, as well as forces produced by cell migration.

Understanding the forces responsible for dissociating and unfolding the GAIN domain from the NTF/CTF complex during force loading at a rate of 1 pN/s is of great significance in physiological contexts. Cell migration speeds are typically slow, on the order of nanometers per second [55]. The large ECR of aGPCRs is connected to the first helix of the 7TM domain by a flexible, membrane-proximal peptide linker, and it consists of multiple structural subdomains linked by flexible connectors. These subdomains are approximately *l* ~ 3 – 5 nm in size and can rotate around the flexible connectors, contributing to a force-extension curve that reflects force-dependent orientations of the subdomains along the direction of force. The magnitude of forces required to orient a rigid structure along the direction of force can be estimated by 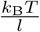, where *k*_B_ is the Boltzmann constant and *T* is the temperature, which falls within the range of a few pN. Therefore, to generate forces of a few pN, the ECR must be extended by at least 10 nm for GPR56 that contains three structral domains (PLL, GAIN A and GAIN B) or longer for larger ECRs of other aGPCRs, resulting in a typical loading rate in the order of pN/s. Thus, during slow cell migration, when the long NTF of an aGPCR adheres stably to the extracellular matrix (ECM), the case with the artificial SpyCatcher-SpyTag linkage that we used in our cell migration assay, the force on the aGPCR is likely to increase at a rate of a few pN per second until all domains are unfolded. Similarly, when the NTF of an aGPCR is associated with a ligand on the plasma membrane of another cell, such as the ligand (CD55)-bound CD97/ADGRE5 [56], membrane deformation induced by cytoskeletal contractions or external forces applied to the cells can potentially generate forces that are transmitted to the GAIN domain, causing its unfolding and/or dissociation.

Notably, unfolding at pN range of forces at loading rates in the order of pN/s is commonly observed for force-bearing structural domains maintained with non-covalent interactions, such as the immunoglobulin (Ig) domain from titin [57], *α*-helical bundles from dystrophin [58], *α*-catenin [59], and talin [52], and the 7TM of GPCR [60]. It is reasonable to hypothesize that the unfolding of the GAIN domain could lead to decryption of the Stachel agonist and facilitate the dissociation of the NTF and CTF subunits [26]. This is consistent with our observation of NTF/CTF dissociation in our cell migration assay, where we used an artificial SpyCatcher-SpyTag linkage.

The proposed mechanical activation mechanism has been suggested to occur through either the force-dependent dissociation of NTF/CTF of self-cleavable aGPCRs or conformational changes in the GAIN domain of non-cleavable aGPCRs. However, until very recently, the required force range and time scale for these processes to occur in any aGPCR remained unknown. A recent study by Zhong et al., which was uploaded to BioRxiv [61] during the preparation of our manuscript, reports on a single-molecule study of the mechanical stability of a different aGPCR (ADGRL3/LPHN3). The authors show that constant forces in the 1-10 pN range are sufficient to dissociate the NTF from CTF within seconds to minutes. However, from that study it remains unclear whether the applied forces could cause GAIN domain conformational changes before dissociation and whether this phenomenon can be extended to other aGPCRs.

Our manuscript addresses these gaps by providing missing information. We demonstrate that at a loading rate of 1 pN/s, GAIN domain unfolding over pN forces is a conserved property for all three aGPCRs tested in this study comparing these mechanical properties across three different aGPCR subfamilies (B, G, and L). Furthermore, extensive GAIN domain unfolding precedes the NTF/CTF dissociation of GPR56 and LPHN1. Our results strongly suggest that the sensitivity of the GAIN domain conformation during physiological stretching processes is a conserved property for aGPCRs.

According to Zhong et al. [61], the GAIN domain of auto-cleaved LPHN3 was found to be remarkably stable and remained undissociated even after several days in solution. This is in stark contrast to the observation that when subjected to pN forces, dissociation of LPHN3’s NTF/CTF occurs within seconds to minutes. The authors of the study speculated that force-dependent dissociation could occur through a different transition pathway, or that additional force-dependent conformational states of the GAIN domain could modulate the dissociation rate. In our study, we found that for both GPR56 and LPHN1, extensive unfolding of the GAIN domain preceded NTF/CTF dissociation. Based on this observation, we postulate that the unfolding of the GAIN domains likely destabilizes the auto-cleaved GAIN domains culminating in NTF/CTF dissociation.

While our study provides insights into the force-dependent unfolding of the GAIN domain and the dissociation of NTF/CTF, it is limited to in vitro experiments. Therefore, the physiological relevance of these findings requires further investigation, as in vivo aGPCRs are subject to more complex dynamic mechanical force environment than those applied in our single-molecule experiments. Additionally, the artificial SpyCatcher ligand used in our cell spreading assay forms a covalent bond with the SpyTag on the NTF of GPR56. However, under physiological conditions the force-transmitting ligand-NTF interface forms through non-covalent interactions and has a limited lifetime. The duration of the force transmission on aGPCRs is determined by the force-dependent lifetime of this interface, which has yet to be determined. Only when the force duration timescale is comparable to or longer than those involved in GAIN domain unfolding and NTF/CTF dissociation can the findings from this study form a basis for interpreting aGPCR-mediated mechanosensing and mechanotransduction. Finally, our observations do not address how the observed GAIN unfolding and NTF/CTF dissociation lead to mechanical activation of aGPCRs, but previous studies have offered complementary approaches to investigate this in the future [62, 63]. Therefore, more comprehensive studies are needed to fully understand the role of mechanical activation in aGPCR signaling, which is the focus of our ongoing research.

In conclusion, this study provides important insights into the mechanical activation hypotheses of aGPCRs [9, 64]. The results suggest that the GAIN domain structural stability and the NTF/CTF dissociation are sensitive to the physiological range of tensile forces. Further investigations are necessary to determine the physiological relevance of these findings and to fully understand the role of mechanical activation in aGPCR signaling. These insights could be useful for the development of drugs that target aGPCRs and for understanding the role of aGPCRs in various physiological processes.

## V. AUTHOR CONTRIBUTIONS

C.F., W.H., Q.T., S.G. performed the experiments; G.S. and J.Y. designed the single-molecule studies. M.N. expressed and purified the protein constructs. C.F., T.L. and J.Y. designed the cell migration assay. G.S. and J.Y. co-supervised the single-molecule studies.

## VI. DECLARATION OF COMPETING INTEREST

The authors declare that they have no known competing financial interests or personal relationships that could have appeared to influence the work reported in this paper.

## VII. ACKNOWLEDGEMENT

J.Y. acknowledges funding by the Singapore Ministry of Education Academic Research Fund Tier 3 (MOE Grant No: MOET32021-0003), and National Research Foundation, Prime Ministers Office, Singapore and the Ministry of Education under the Research Centres of Excellence programme. T.L. acknowledges funding by the Deutsche Forschungsgemeinschaft through SFB1423, project number 421152132, subproject B06. G. S. acknowledges funding by the National Nature Science Foundation of China Grant 31770898.

## VIII. SUPPLEMENTARY INFORMATION

The supplementary information describes: 1) derivation of Eq. 1 in the main text (S1); 2) the numerical solution of the Fokker-Planck equation (S2); 3) the conformation free energy of a molecule stretched by an external constant force (S3)).

## Notes

### Competing Interest Statement

The authors have declared no competing interest.

## References

[1] D. Hilger, M. Masureel, and B. K. Kobilka, Structure and dynamics of gpcr signaling complexes, Nature structural & molecular biology 25, 4 (2018).

[2] R. H. Purcell and R. A. Hall, Adhesion g protein–coupled receptors as drug targets, Annual review of pharmacology and toxicology 58, 429 (2018).

[3] O. Civelli, R. K. Reinscheid, Y. Zhang, Z. Wang, R. Fredriksson, and H. B. Schiöth, G protein–coupled receptor deorphanizations, Annual review of pharmacology and toxicology 53, 127 (2013).

[4] M. C. Lagerström and H. B. Schiöth, Structural diversity of g protein-coupled receptors and significance for drug discovery, Nature reviews Drug discovery 7, 339 (2008).

[5] H. B. Schiöth and R. Fredriksson, The grafs classification system of g-protein coupled receptors in comparative perspective, General and comparative endocrinology 142, 94 (2005).

[6] D. Araç, N. Sträter, and E. Seiradake, Understanding the structural basis of adhesion gpcr functions, in Adhesion G protein-coupled receptors (Springer, 2016) pp. 67–82.

[7] A. J. McKnight and S. Gordon, The egf-tm7 family: unusual structures at the leukocyte surface, Journal of leukocyte biology 63, 271 (1998).

[8] J. Hamann, G. Aust, D. Araç, F. B. Engel, C. Formstone, R. Fredriksson, R. A. Hall, B. L. Harty, C. Kirchhoff, B. Knapp, et al., International union of basic and clinical pharmacology. xciv. adhesion g protein–coupled receptors, Pharmacological reviews 67, 338 (2015).

[9] A. Vizurraga, R. Adhikari, J. Yeung, M. Yu, and G. G. Tall, Mechanisms of adhesion g protein–coupled receptor activation, Journal of Biological Chemistry 295, 14065 (2020).

[10] S. Prömel, M. Frickenhaus, S. Hughes, L. Mestek, D. Staunton, A. Woollard, I. Vakonakis, T. Schöneberg, R. Schnabel, A. P. Russ, et al., The gps motif is a molecular switch for bimodal activities of adhesion class g protein-coupled receptors, Cell reports 2, 321 (2012).

[11] D. Araç, A. A. Boucard, M. F. Bolliger, J. Nguyen, S. M. Soltis, T. C. Südhof, and A. T. Brunger, A novel evolutionarily conserved domain of cell-adhesion gpcrs mediates autoproteolysis, The EMBO journal 31, 1364 (2012).

[12] H. M. Stoveken, A. G. Hajduczok, L. Xu, and G. G. Tall, Adhesion g protein-coupled receptors are activated by exposure of a cryptic tethered agonist, Proceedings of the National Academy of Sciences 112, 6194 (2015).

[13] I. Liebscher, J. Schön, S. C. Petersen, L. Fischer, N. Auerbach, L. M. Demberg, A. Mogha, M. Cöster, K.-U. Simon, S. Rothemund, et al., A tethered agonist within the ectodomain activates the adhesion g protein-coupled receptors gpr126 and gpr133, Cell reports 9, 2018 (2014).

[14] J. X. Gray, M. Haino, M. J. Roth, J. E. Maguire, P. N. Jensen, A. Yarme, M.-A. Stetler-Stevenson, U. Siebenlist, and K. Kelly, Cd97 is a processed, seven-transmembrane, heterodimeric receptor associated with inflammation., Journal of immunology 157, 5438 (1996).

[15] V. G. Krasnoperov, M. A. Bittner, R. Beavis, Y. Kuang, K. V. Salnikow, O. G. Chepurny, A. R. Little, A. N. Plotnikov, D. Wu, R. W. Holz, et al., α-latrotoxin stimulates exocytosis by the interaction with a neuronal g-protein-coupled receptor, Neuron 18, 925 (1997).

[16] H.-H. Lin, G.-W. Chang, J. Q. Davies, M. Stacey, J. Harris, and S. Gordon, Autocatalytic cleavage of the emr2 receptor occurs at a conserved g protein-coupled receptor proteolytic site motif, Journal of Biological Chemistry 279, 31823 (2004).

[17] G. Beliu, S. Altrichter, R. Guixà-González, M. Hemberger, I. Brauer, A.-K. Dahse, N. Scholz, R. Wieduwild, A. Kuhlemann, H. Batebi, et al., Tethered agonist exposure in intact adhesion/class b2 gpcrs through intrinsic structural flexibility of the gain domain, Molecular cell 81, 905 (2021).

[18] T. Langenhan, Adhesion g protein–coupled receptors— candidate metabotropic mechanosensors and novel drug targets, Basic & clinical pharmacology & toxicology 126, 5 (2020).

[19] T. Langenhan, X. Piao, and K. R. Monk, Adhesion g protein-coupled receptors in nervous system development and disease, Nature Reviews Neuroscience 17, 550 (2016).

[20] N. Scholz, Cancer cell mechanics: adhesion g protein-coupled receptors in action?, Frontiers in oncology 8, 59 (2018).

[21] Z. Kan, B. S. Jaiswal, J. Stinson, V. Janakiraman, D. Bhatt, H. M. Stern, P. Yue, P. M. Haverty, R. Bourgon, J. Zheng, et al., Diverse somatic mutation patterns and pathway alterations in human cancers, Nature 466, 869 (2010).

[22] C. J. Folts, S. Giera, T. Li, and X. Piao, Adhesion g protein-coupled receptors as drug targets for neurological diseases, Trends in pharmacological sciences 40, 278 (2019).

[23] H.-H. Lin, C.-C. Hsiao, C. Pabst, J. Hébert, T. Schöneberg, and J. Hamann, Adhesion gpcrs in regulating immune responses and inflammation, Advances in immunology 136, 163 (2017).

[24] D. Liu, L. Duan, L. B. Rodda, E. Lu, Y. Xu, J. An, L. Qiu, F. Liu, M. R. Looney, Z. Yang, et al., Cd97 promotes spleen dendritic cell homeostasis through the mechanosensing of red blood cells, Science 375, eabi5965 (2022).

[25] J. Yeung, R. Adili, E. N. Stringham, R. Luo, A. Vizurraga, L. K. Rosselli-Murai, H. M. Stoveken, M. Yu, X. Piao, M. Holinstat, et al., Gpr56/adgrg1 is a platelet collagen-responsive gpcr and hemostatic sensor of shear force, Proceedings of the National Academy of Sciences 117, 28275 (2020).

[26] N. Scholz, A.-K. Dahse, M. Kemkemer, A. Bormann, G. M. Auger, F. Vieira Contreras, L. F. Ernst, H. Staake, M. B. Körner, M. Buhlan, et al., Molecular sensing of mechano-and ligand-dependent adhesion gpcr dissociation, Nature, Epub ahead of print. 10.1038/s41586 (2023).

[27] N. Scholz, C. Guan, M. Nieberler, A. Grotemeyer, I. Maiellaro, S. Gao, S. Beck, M. Pawlak, M. Sauer, E. Asan, et al., Mechano-dependent signaling by lat-rophilin/cirl quenches camp in proprioceptive neurons, Elife 6, e28360 (2017).

[28] R. Sando, X. Jiang, and T. C. Südhof, Latrophilin gpcrs direct synapse specificity by coincident binding of flrts and teneurins, Science 363, eaav7969 (2019).

[29] X. Barros-Álvarez, R. M. Nwokonko, A. Vizurraga, D. Matzov, F. He, M. M. Papasergi-Scott, M. J. Robertson, O. Panova, E. H. Yardeni, A. B. Seven, et al., The tethered peptide activation mechanism of adhesion gpcrs, Nature 604, 757 (2022).

[30] P. Xiao, S. Guo, X. Wen, Q.-T. He, H. Lin, S.-M. Huang, L. Gou, C. Zhang, Z. Yang, Y.-N. Zhong, et al., Tethered peptide activation mechanism of the adhesion gpcrs adgrg2 and adgrg4, Nature 604, 771 (2022).

[31] X. Qu, N. Qiu, M. Wang, B. Zhang, J. Du, Z. Zhong, W. Xu, X. Chu, L. Ma, C. Yi, et al., Structural basis of tethered agonism of the adhesion gpcrs adgrd1 and adgrf1, Nature 604, 779 (2022).

[32] Y.-Q. Ping, P. Xiao, F. Yang, R.-J. Zhao, S.-C. Guo, X. Yan, X. Wu, C. Zhang, Y. Lu, F. Zhao, et al., Structural basis for the tethered peptide activation of adhesion gpcrs, Nature 604, 763 (2022).

[33] H. Lin, P. Xiao, R.-Q. Bu, S. Guo, Z. Yang, D. Yuan, Z.-L. Zhu, C.-X. Zhang, Q.-T. He, C. Zhang, et al., Structures of the adgrg2–gs complex in apo and ligand-bound forms, Nature Chemical Biology 18, 1196 (2022).

[34] Y. Qian, Z. Ma, C. Liu, X. Li, X. Zhu, N. Wang, Z. Xu, R. Xia, J. Liang, Y. Duan, et al., Structural insights into adhesion gpcr adgrl3 activation and gq, gs, gi, and g12 coupling, Molecular Cell 82, 4340 (2022).

[35] X. Zhu, Y. Qian, X. Li, Z. Xu, R. Xia, N. Wang, J. Liang, H. Yin, A. Zhang, C. Guo, et al., Structural basis of adhesion gpcr gpr110 activation by stalk peptide and g-proteins coupling, Nature Communications 13, 5513 (2022).

[36] S. C. Petersen, R. Luo, I. Liebscher, S. Giera, S.-J. Jeong, A. Mogha, M. Ghidinelli, M. L. Feltri, T. Schöneberg, X. Piao, et al., The adhesion gpcr gpr126 has distinct, domain-dependent functions in schwann cell development mediated by interaction with laminin-211, Neuron 85, 755 (2015).

[37] N. Scholz, J. Gehring, C. Guan, D. Ljaschenko, R. Fischer, V. Lakshmanan, R. J. Kittel, and T. Langenhan, The adhesion gpcr latrophilin/cirl shapes mechanosensation, Cell reports 11, 866 (2015).

[38] S. E. Boyden, A. Desai, G. Cruse, M. L. Young, H. C. Bolan, L. M. Scott, A. R. Eisch, R. D. Long, C.-C. R. Lee, C. L. Satorius, et al., Vibratory urticaria associated with a missense variant in adgre2, New England Journal of Medicine 374, 656 (2016).

[39] O. N. Karpus, H. Veninga, R. M. Hoek, D. Flierman, J. D. van Buul, C. C. Vandenakker, E. vanBavel, M. E. Medof, R. A. van Lier, K. A. Reedquist, et al., Shear stress–dependent downregulation of the adhesion-g protein–coupled receptor cd97 on circulating leukocytes upon contact with its ligand cd55, The Journal of Immunology 190, 3740 (2013).

[40] J. P. White, C. D. Wrann, R. R. Rao, S. K. Nair, M. P. Jedrychowski, J.-S. You, V. Martínez-Redondo, S. P. Gygi, J. L. Ruas, T. A. Hornberger, et al., G protein-coupled receptor 56 regulates mechanical overload-induced muscle hypertrophy, Proceedings of the National Academy of Sciences 111, 15756 (2014).

[41] C. Wilde, L. Fischer, V. Lede, J. Kirchberger, S. Rothemund, T. Schöneberg, and I. Liebscher, The constitutive activity of the adhesion gpcr gpr114/adgrg5 is mediated by its tethered agonist, The FASEB Journal 30, 666 (2016).

[42] I. Liebscher and T. Schöneberg, Tethered agonism: a common activation mechanism of adhesion gpcrs, in Adhesion G Protein-coupled Receptors (Springer, 2016) pp. 111–125.

[43] A. Kishore and R. A. Hall, Versatile signaling activity of adhesion gpcrs, in Adhesion G Protein-coupled Receptors (Springer, 2016) pp. 127–146.

[44] M. Nieberler, R. J. Kittel, A. G. Petrenko, H.-H. Lin, and T. Langenhan, Control of adhesion gpcr function through proteolytic processing, in Adhesion G Protein-coupled Receptors (Springer, 2016) pp. 83–109.

[45] S. M. Sigoillot, K. R. Monk, X. Piao, F. Selimi, and B. L. Harty, Adhesion gpcrs as novel actors in neural and glial cell functions: from synaptogenesis to myelination, in Adhesion G Protein-coupled Receptors (Springer, 2016) pp. 275–298.

[46] P. Roca-Cusachs, V. Conte, and X. Trepat, Quantifying forces in cell biology, Nature cell biology 19, 742 (2017).

[47] M. Huse, Mechanical forces in the immune system, Nature Reviews Immunology 17, 679 (2017).

[48] H. Chen, H. Fu, X. Zhu, P. Cong, F. Nakamura, and J. Yan, Improved high-force magnetic tweezers for stretching and refolding of proteins and short dna, Biophysical journal 100, 517 (2011).

[49] X. Zhao, X. Zeng, C. Lu, and J. Yan, Studying the mechanical responses of proteins using magnetic tweezers, Nanotechnology 28, 414002 (2017).

[50] G. S. Salzman, S. D. Ackerman, C. Ding, A. Koide, K. Leon, R. Luo, H. M. Stoveken, C. G. Fernandez, G. G. Tall, X. Piao, et al., Structural basis for regulation of gpr56/adgrg1 by its alternatively spliced extracellular domains, Neuron 91, 1292 (2016).

[51] B. Zakeri, J. O. Fierer, E. Celik, E. C. Chittock, U. Schwarz-Linek, V. T. Moy, and M. Howarth, Peptide tag forming a rapid covalent bond to a protein, through engineering a bacterial adhesin, Proceedings of the National Academy of Sciences 109, E690 (2012).

[52] M. Yao, B. T. Goult, B. Klapholz, X. Hu, C. P. Toseland, Y. Guo, P. Cong, M. P. Sheetz, and J. Yan, The mechanical response of talin, Nature communications 7, 11966 (2016).

[53] A. Kishore, R. H. Purcell, Z. Nassiri-Toosi, and R. A. Hall, Stalk-dependent and stalk-independent signaling by the adhesion g protein-coupled receptors gpr56 (adgrg1) and bai1 (adgrb1), Journal of Biological Chemistry 291, 3385 (2016).

[54] H. Chen, F. Nakamura, H. Fu, P. Cong, M. P. Sheetz, and J. Yan, Unfolding and refolding dynamics of filamin a protein under constant forces, Biophysical Journal 98, 753a (2010).

[55] L. Ge, L. Yang, R. Bron, J. K. Burgess, and P. van Rijn, Topography-mediated fibroblast cell migration is influenced by direction, wavelength, and amplitude, ACS Applied Bio Materials 3, 2104 (2020).

[56] M. Niu, S. Xu, J. Yang, D. Yao, N. Li, J. Yan, G. Zhong, and G. Song, Structural basis for cd97 recognition of the decay-accelerating factor cd55 suggests mechanosensitive activation of adhesion gpcrs, Journal of Biological Chemistry 296 (2021).

[57] M. Yu, J.-H. Lu, S. Le, and J. Yan, Unexpected low mechanical stability of titin i27 domain at physiologically relevant temperature, The Journal of Physical Chemistry Letters 12, 7914 (2021).

[58] S. Le, M. Yu, L. Hovan, Z. Zhao, J. Ervasti, and J. Yan, Dystrophin as a molecular shock absorber, ACS nano 12, 12140 (2018).

[59] M. Yao, W. Qiu, R. Liu, A. K. Efremov, P. Cong, R. Seddiki, M. Payre, C. T. Lim, B. Ladoux, R.-M. Mege, et al., Force-dependent conformational switch of *α*-catenin controls vinculin binding, Nature communications 5, 4525 (2014).

[60] H.-K. Choi, D. Min, H. Kang, M. J. Shon, S.-H. Rah, H. C. Kim, H. Jeong, H.-J. Choi, J. U. Bowie, and T.-Y. Yoon, Watching helical membrane proteins fold reveals a common n-to-c-terminal folding pathway, Science 366, 1150 (2019).

[61] B. L. Zhong, C. E. Lee, V. T. Vachharajani, T. C. Sudhof, and A. R. Dunn, Piconewton forces mediate gain domain dissociation of the latrophilin-3 adhesion gpcr, bioRxiv, 2023 (2023).

[62] G. Stephan, J. D. Frenster, I. Liebscher, and D. G. Placantonakis, Activation of the adhesion g protein–coupled receptor gpr133 by antibodies targeting its n-terminus, Journal of Biological Chemistry 298 (2022).

[63] J. Mitgau, J. Franke, C. Schinner, G. Stephan, S. Berndt, D. G. Placantonakis, H. Kalwa, V. Spindler, C. Wilde, and I. Liebscher, The n terminus of adhesion g protein– coupled receptor gpr126/adgrg6 as allosteric force integrator, Frontiers in Cell and Developmental Biology, 1169 (2022).

[64] N. Scholz, K. R. Monk, R. J. Kittel, and T. Langenhan, Adhesion gpcrs as a putative class of metabotropic mechanosensors, Adhesion G protein-coupled receptors: molecular, physiological and pharmacological principles in health and disease, 221 (2016).

